# Patterns of Ongoing Thought in the Real-World

**DOI:** 10.1101/2022.10.05.510994

**Authors:** Bridget Mulholland, Ian Goodall-Halliwell, Raven Wallace, Louis Chitiz, Brontë Mckeown, Aryanna Rastan, Giulia Poerio, Robert Leech, Adam Turnbull, Arno Klein, Wil Van Auken, Michael Milham, Jeffrey Wammes, Elizabeth Jefferies, Jonathan Smallwood

**Author notes:** Corresponding author (BM).

## Abstract

Previous research has indicated that health and well-being are impacted on by both the way we think, and the things we do. In the laboratory, studies suggest that specific task contexts affect this process because the people we are with, the places we are in, and the activities we perform may influence our thought patterns. In our study participants completed multi-dimensional experience-sampling surveys eight times per day for 5 days to generate thought data across a variety of dimensions in daily life. Principal component analysis was used to decompose the experience sampling data, and linear mixed modelling related these patterns to the activity in daily life in which they emerged. Our study replicated the influence of socializing on patterns of ongoing thought observed in our prior study and established that this is part of a broader set of relationship that links our current activities to how our thoughts are organised in daily life. We also found that factors such as time of day and the physical location are associated with reported patterns of thought, factors that are important for future studies to explore. Our study suggests that sampling thinking in the real world may be able to provide a set of comprehensive thinking-activity mappings that will be useful to researchers and health care professionals interested in health and well-being.

## Introduction

A core goal of cognitive science is to understand the processes that support cognition and contemporary work suggests that the content and form of the way we think varies widely across people, places, and activities (Smallwood et al., 2021). Variation in how we think and feel is well known to contribute to health and well-being (Fitzgerald et al., 2008), and the sorts of activities we engage in are also important (Ingram et al., 2020). Consequently, research is needed to determine how thought emerges across these different contexts, particularly within natural environments. This will help build better connections between theoretical models of how we think, and how these will play out in the activities we perform in daily life. One aim of our study, therefore, was to map ongoing patterns of thought and behaviour in a real-world context in order to provide a preliminary description of how thoughts map onto activities in daily life.

Important aspects of cognition can be measured in lab-based settings, allowing insight into processes underlying human thought. However, it is difficult to gain similar information in daily life (Kingstone et al., 2003). As noted in Kingstone et al. (2003), research based in natural environments is needed to establish ecological validity within real-world contexts. Consistent with this perspective, previous research suggests that lab-based descriptions of ongoing thought may not generalize to real-world contexts (Ho et al., 2020). Accordingly, it would be useful to gain contextualised measurements of thinking in the real-world to provide a provisional description of the factors that impact the landscape of thinking in daily life.

Our study set out to use the technique of experience-sampling (ES) to provide a description of thinking in daily life. ES allows researchers to capture what people are thinking during everyday activities and lab-based tasks (Conner et al., 2009; Smallwood et al., 2021). This technique has been used to provide descriptions of psychopathology (Myin-Germeys et al., 2018) and emotion in the real-world (Zelenski & Larsen, 2000). Studies have also examined how states like mind-wandering emerge in daily life (Franklin et al., 2013; Kane et al., 2007, 2017).

Our current study sought to extend these approaches via the use of multi-dimensional experience-sampling (MDES) to map patterns of ongoing thought onto primary activities in both real-world and laboratory settings (Smallwood et al., 2016; Ho et al., 2020). MDES asks participants to describe their thinking across several dimensions (Smallwood et al., 2016). For example, across a “Task” dimension, participants might be asked to score themselves on a 1 to 5 Likert scale (1 = Not at all, 5 = Completely) in relation to the associated statement, “My thoughts were focused on the task I was performing” (Smallwood et al., 2016). MDES questions are traditionally decomposed via principal component analysis (PCA) into a low dimensional space, and these dimensions can be visualised as word clouds (see Figure 4). MDES is a powerful technique that can be used to determine associations between patterns of thought, making links to brain activity (Konu et al., 2020; Turnbull et al., 2019), and in the lab can be linked to states like autism (Turnbull et al., 2020) and attention deficit disorder (Vatansever et al., 2019).

Recently, Mckeown et al. (2021) used MDES to map ongoing thought in the real-world onto primary activities during the first coronavirus disease 2019 (COVID-19) lockdown in the United Kingdom. They found specific behavioural changes resultant of lockdown-reduced opportunities for working and socializing, leading to unique changes in patterns of ongoing thought. One goal of this study was to replicate the influence of socializing on patterns of ongoing thought. We hypothesized that episodic social cognition thought patterns, which relate to thinking about other people, would dominate activities that involved other people, as seen in Mckeown et al. (2021).

As well as replicating this prior study, we also hoped to understand how the activity that a person is performing in daily life is reflected in their thought patterns, as captured by MDES. In the laboratory, Konu et al. (2021) used MDES to investigate the influence of a task on patterns of ongoing thought in lab-based settings via task mapping. They discovered that thought patterns differ under specific task contexts in lab-based settings. Specifically, episodic social cognition though patterns were present when tasks involved thinking about friends and were absent when watching affective TV clips and engaging in memory tasks (Konu et al., 2021). In contrast, patterns of detailed task focused thoughts were most common when performing tasks that depended on executive control (working memory, task switching). Thus, a second goal of this study was to extend research to examine more generally whether activities in the real-world impact on a person’s thinking.

We also had two more exploratory aims. Studies have suggested that whether a person is indoors or outdoors can impact on their psychological state (Duvall, 2011; Weng & Chiang, 2014). Since natural variation in where participants where when the probe occurred would allow us to sample thinking in a variety of different locations in our study, we also ascertained whether the participants were indoors or outdoors when the probe occurred. Using this data, we explored whether this impacted on their experience. Finally, certain types of activities are more likely to occur at certain times of the day. As a final exploratory goal, therefore, we examined whether the time of the day in which the MDES probe occurred was reflected in the patterns of thought the participants described.

In summary, the broad goal of our study was to examine how thinking patterns in the real-world relate to the activity in which the experience occurred. Based on prior work we expected to replicate the influence of socializing on patterns of ongoing thought (Mckeown et al., 2021). Second, we aimed to determine whether there is a relationship between activities and ongoing thought patterns in the real-world that parallels those seen in tasks (Konu et al., 2021). Third, we aimed to discover whether MDES was linked to variation in time of day or location.

## Method

### Participants

This study was granted initial ethics clearance by the Queen’s University Health Sciences & Affiliated Teaching Hospitals Research Ethics Board (HSREB). Participants were recruited between February 2022 and April 2022 though the Queen’s University Psychology Participant Pool. This recruitment timeline was determined by the Psychology Participant Pool participation end date. Eligible participants were Queen’s University students enrolled in designated first- and second-year psychology courses. Participants gave informed, written consent via electronic documentation prior to taking part in any research activities. Participants were awarded 2.0 course credits and also fully debriefed upon the completion of this study. A total of 101 participants (women = 82, men = 13, non-binary = 2, did not specify = 3; age: M = 21.11; SD = 5.33; and range = 18 to 52) completed MDES surveys with additional stress, environment, location, and activity questions.

### Procedure

Participants were emailed a MindLogger Pilot invitation for an applet called “THOUGHTLOG,” which they were instructed to accept. MindLogger Pilot is a smartphone application that allows researchers to collect, analyze, and visualize data through custom activities such as surveys, quizzes, digital diaries, and cognitive tasks, using mobile devices (Child Mind Institute, 2022). The THOUGHTLOG applet contained an MDES survey with additional stress, environment, location, and activity questions that participants completed for this study. Participants were required to download the MindLogger Pilot application onto their smartphone to access the THOUGHTLOG applet, and consequently, the MDES survey and additional questions. Participants were notified to complete the THOUGHTLOG applet by the MindLogger Pilot application eight times daily for 5 days between the hours of 7:00 AM and 11:00 PM. Each prompt was delivered within a specific 2-hour time interval. The MDES survey and additional questions were not accessible outside of their associated 2-hour time interval.

### Multi-Dimensional Experience-Sampling and Additional Questions

Participants were asked 14 multi-dimensional experience-sampling (MDES) questions about the content of their thoughts immediately before being notified by MindLogger Pilot across a variety of dimensions (Table 1). Participants were also asked to rate their stress level immediately before being notified by MindLogger Pilot. Next, participants were asked questions about their social environments immediately before being notified by MindLogger Pilot (Table 2). Additionally, participants were asked to indicate their type of location and primary activity immediately before being notified by MindLogger Pilot (Table 3, 4). The primary activity list was developed from the day reconstruction method (Kahneman et al., 2004) and modified based on the activity options in Mckeown et al. (2021).

**Table 1.**
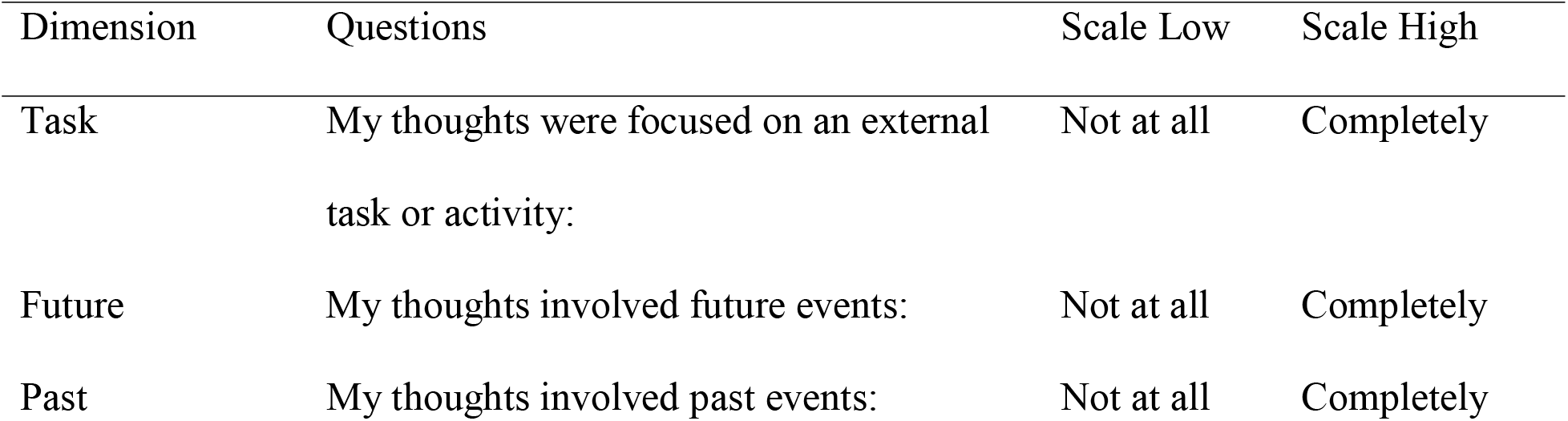
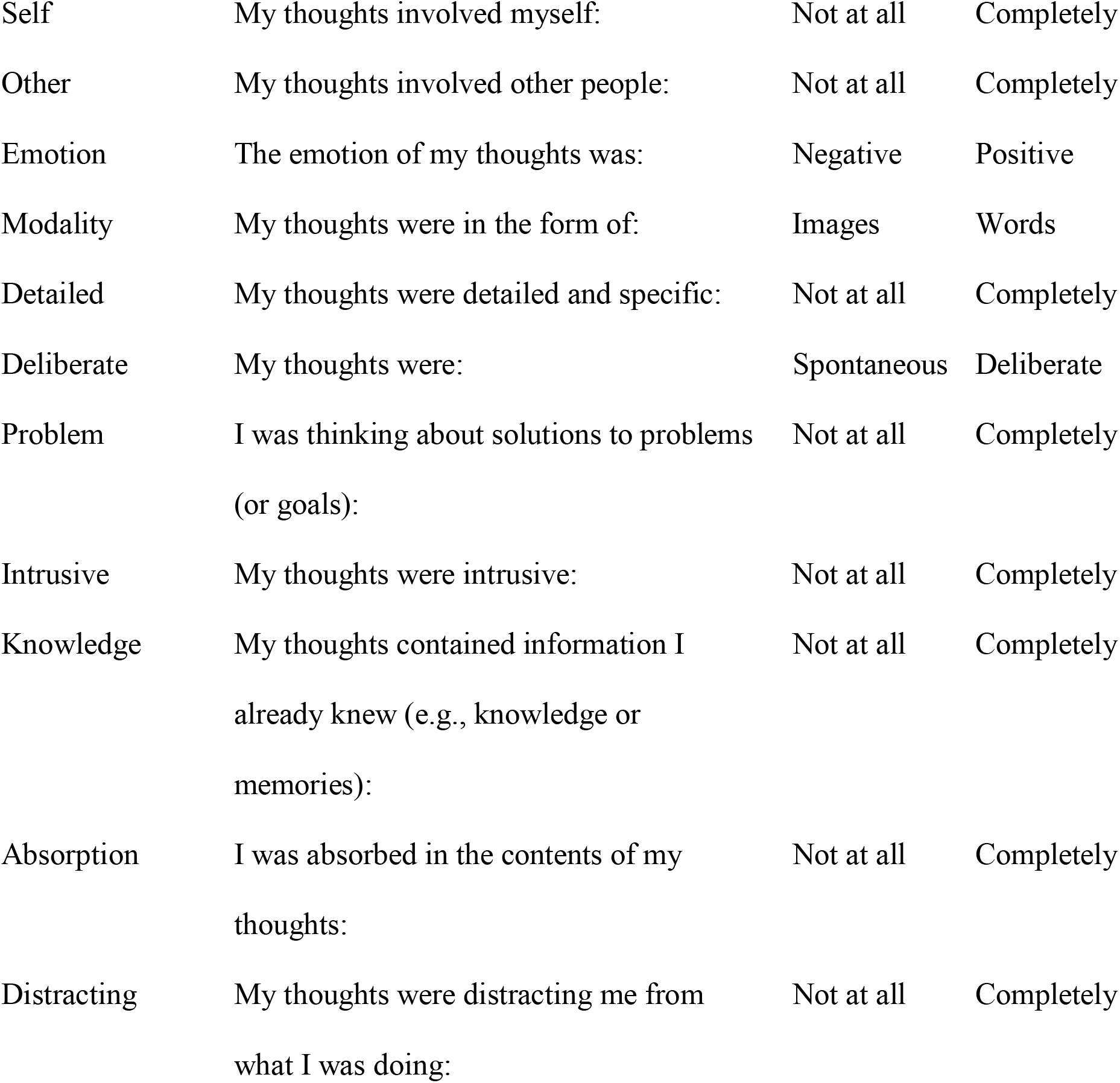
Summary of the multi-dimensional experience-sampling (MDES) questions. Participants rated statements on a 1 to 5 Likert scale.

| Dimension | Questions | Scale Low | Scale High |
| --- | --- | --- | --- |
| Task | My thoughts were focused on an external task or activity: | Not at all | Completely |
| Future | My thoughts involved future events: | Not at all | Completely |
| Past | My thoughts involved past events: | Not at all | Completely |
| Self | My thoughts involved myself: | Not at all | Completely |
| Other | My thoughts involved other people: | Not at all | Completely |
| Emotion | The emotion of my thoughts was: | Negative | Positive |
| Modality | My thoughts were in the form of: | Images | Words |
| Detailed | My thoughts were detailed and specific: | Not at all | Completely |
| Deliberate | My thoughts were: | Spontaneous | Deliberate |
| Problem | I was thinking about solutions to problems<br>(or goals): | Not at all | Completely |
| Intrusive | My thoughts were intrusive: | Not at all | Completely |
| Knowledge | My thoughts contained information I<br>already knew (e.g., knowledge or<br>memories): | Not at all | Completely |
| Absorption | I was absorbed in the contents of my<br>thoughts: | Not at all | Completely |
| Distracting | My thoughts were distracting me from<br>what I was doing: | Not at all | Completely |

**Table 2.** Summary of the social environment questions.

| Environment | Question | Environment List |
| --- | --- | --- |
| Physical | Where you alone, or | Alone |
|  | physically with other | Around people but not interacting with them |
|  | people? | Around people and interacting with them |
| Virtual | Where you alone, or | Alone |
|  | virtually with other | Around people but not interacting with them |
|  | people? | (e.g., reading messages but not replying,<br>being on a video call but not participating,<br>etc.) |
|  |  | Around people and interacting with them<br>(e.g., direct communication with another<br>person by text, instant messaging, calling, or<br>video calling, etc.) |

**Table 3.** Summary of the location questions. If participants selected “Inside (other),” or “Outside (other),” they were asked to specify their location.

| Question | Location List |
| --- | --- |
| Where were you? | Inside a home |
|  | Inside a shop |
|  | Inside a workplace |
|  | Inside (other) |
|  | Outside in a city or town |
|  | Outside in nature |
|  | Outside (other) |

**Table 4.** Summary of the primary activity questions. If participants selected “Other activity,” they were asked to specify their primary activity.

| Question | Possible Activity Answers |
| --- | --- |
| What were you doing? | Eating |
|  | Homework |
|  | Household chores |
|  | Listening to music |
|  | Napping or resting |
|  | Nothing or waiting |
|  | Personal exercise |
|  | Personal hygiene care |
|  | Physical leisure or sports |
|  | Reading |
|  | Shopping |
|  | Talking in person |
|  | Talking on the phone |
|  | Texting by phone |
|  | Traveling or commuting |
|  | Using a computer or an electronic device |
|  | Walking the dog |
|  | Watching TV |
|  | Working (paid or volunteer) |
|  | Other activity |

### Analysis

#### Code availability statement

All custom code used to prepare data for analysis and figure development is available at https://github.com/ThinCLabQueens.

#### Principal component analysis (PCA)

Common patterns of thought were identified by applying PCA with varimax rotation to all thought data generated from responses to the 14 multi-dimensional experience-sampling (MDES) questions (Table 1) using IBM SPSS (version 28). This is the standard method, as seen in Mckeown et al. (2021) and Konu et al. (2021). The loadings from the four components with an eigenvalue >1 were saved for the generation of thought word clouds. These components were also saved for later use as outcome variables for linear mixed modelling (LMM).

#### Component Reliability

Component reliability analysis was conducted in IBM SPSS (version 28). First, all thought data was randomly shuffled, and then divided into two halves, with each half containing a sample of 729 probes. Next, a column titled “subset” was added to the datasheet. Each row was labelled as “1” or “2” to indicate subset. To assess component reliability, PCA with varimax rotation was utilized, with subset included as a selection variable. Further, factor scores were estimated using the Thurstone regression method for all thought date based on the factors generated from each subset. Afterwards, Pearson correlations were run on the factor scores between each of the factors generated from each subset.

#### Mixed Modelling (LMM), Primary Activity Data

To analyze contextual distributions of thought in relation to activities, LMM was applied to each of the component loadings generated by PCA. Activities were included as a condition of interest, and participants were be included as a random factor. This is the standard method, as seen in Mckeown et al. (2021), and Konu et al. (2021). LMM loadings were saved for the eventual generation of activity word clouds (Figure 4). This analysis is identical to that found in Konu et al. (2021), with the only exception being the use of activities found in the real-world, rather than lab-based tasks.

#### Activity Time Dependence

Analysis of activity time was assessed using SPSS version 28. The “time” variable was recoded into bins that divided the 24-hour period into 6 time bins using a visual binning function. Categorization of bins included early morning (00:00:00 - 10:26:40), late morning (10:33:20 - 12:26:40), early afternoon (12:33:20 - 15:06:40), late afternoon (15:13:20 - 17:40:00), evening (17:46:40 - 20:26:40), and night (20:33:20 - 23:53:20). A frequency analysis was applied to each time bin to evaluate the frequency of reported activities engaged in by participants.

#### Linear Mixed Modelling (LMM), Time of Day Data

To analyzes contextual distributions of thought in relation to time of day, LMM was applied to each of the component loadings generated previously by PCA. Time was included as a condition of interest, and participants were included as a random factor.

## Results

### Patterns of Ongoing Thought

First, mean dimension scores from the thought date were calculated and displayed as a horizontal bar graph (Figure 1A). Next, thought data was decomposed using PCA. The associated scree plot generated from PCA indicated a 4-factor solution, determined by an eigenvalue >1 (Figure 1B). PCA loadings (Table 5) from the four components were used to generate thought word clouds (Figure 1C-F). Thought word clouds were labelled based on MDES dimensions that dominated their composition. Component 1 was labelled “detailed task focus” because of significant loadings for “Detailed,” and “Task” (Figure 1C). Component 2 was labelled “negative intrusive distracting” because of significant loadings for “Emotion,” “Intrusive,” and “Distracting” (Figure 1D). Component 3 was labelled “future problem-solving” because of significant loadings for “Future” and “Problem” (Figure 1E). Component 4 was labelled “episodic social cognition” because of significant loadings for “Past,” “Knowledge,” and “Person” (Figure 1F). Please note that these terms are used for convenience when discussing these patterns, they do not constitute the only label which could be applied to these patterns.

**Figure 1.**
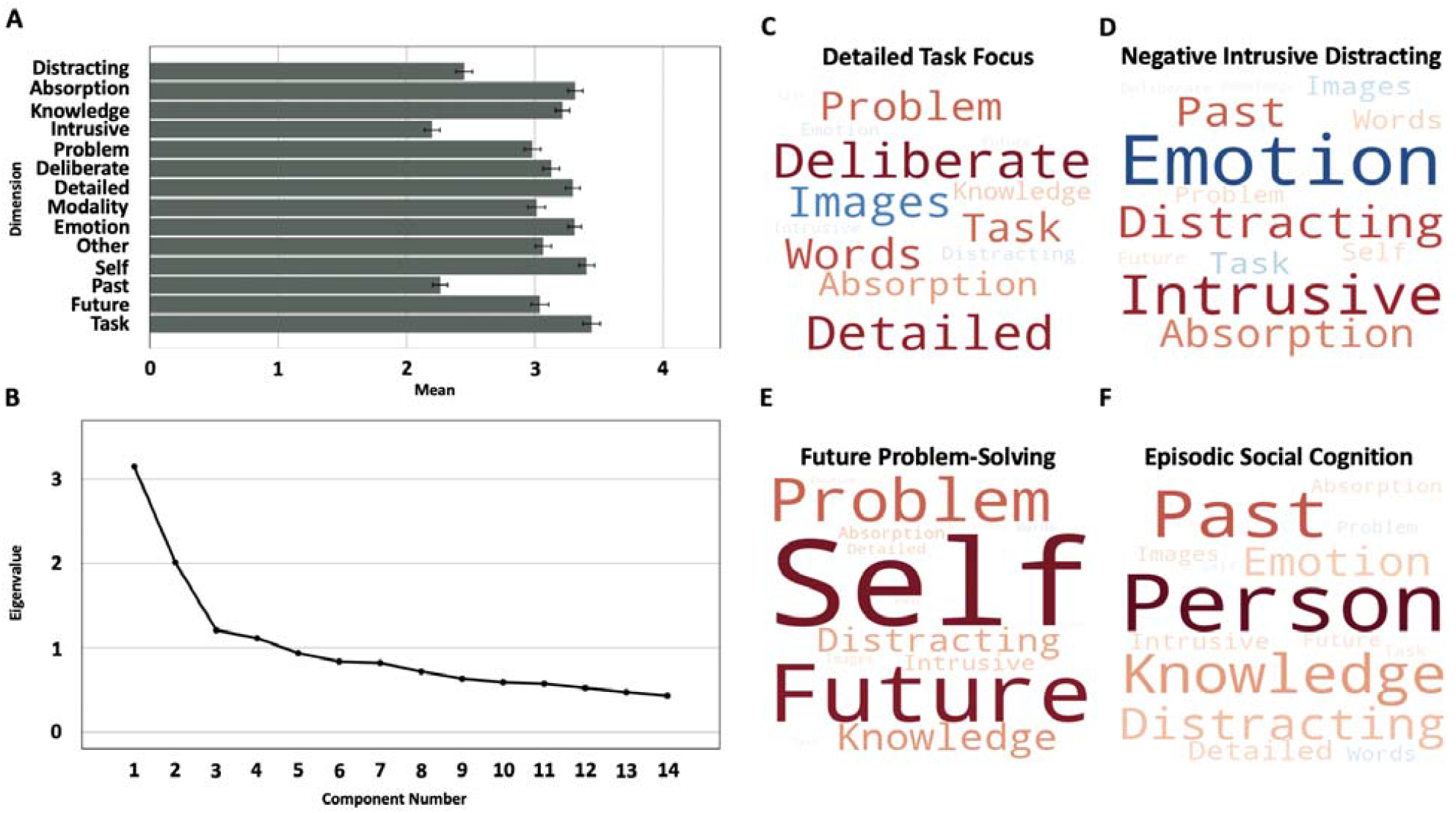
Patterns of ongoing thought identified through 14 multi-dimensional experience-sampling (MDES) probes. (A) Horizontal bar graph of mean dimension scores. Error bars represent 99% CIs. (B) Scree plot generated from PCA of MDES probe data. (C-F) Words represent experience-sampling items, and primary activities. Larger fonts are items with more importance and colour described direction (warm colours relate to positive loadings). (C) Detailed task focus thought word cloud. (D) Negative intrusive distracting thought word cloud. (E) Future problem-solving word cloud. (F) Episodic social cognition thought word cloud.

**Table 5.** Thought data loadings generated by Principal Component Analysis (PCA).

| Dimension | Component |  |  |  |
| --- | --- | --- | --- | --- |
|  | 1 | 2 | 3 | 4 |
| Task | 0.491 | -0.258 | 0.077 | 0.075 |
| Future | 0.054 | 0.079 | 0.763 | 0.124 |
| Past | -0.003 | 0.507 | 0.086 | 0.549 |
| Self | 0.041 | 0.157 | 0.787 | -0.056 |
| Other | -0.001 | 0.019 | -0.021 | 0.845 |
| Emotion | -0.102 | -0.757 | 0.081 | 0.209 |
| Modality | 0.569 | 0.195 | -0.109 | -0.149 |
| Detailed | 0.721 | 0.013 | 0.187 | 0.190 |
| Deliberate | 0.765 | -0.039 | 0.061 | -0.018 |
| Problem | 0.505 | 0.154 | 0.517 | -0.096 |
| Intrusive | -0.073 | 0.718 | 0.221 | 0.177 |
| Knowledge | 0.219 | 0.037 | 0.386 | 0.359 |
| Absorption | 0.370 | 0.424 | 0.223 | 0.135 |
| Distracting | -0.127 | 0.591 | 0.338 | 0.254 |

**Table 6.** Activity data loadings generated by Linear Mixed Modelling (LMM).

| Activity | Component |  |  |  |
| --- | --- | --- | --- | --- |
|  | 1 | 2 | 3 | 4 |
| Eating | -0.194 | -0.04 | 0.15 | 0.126 |
| Homework | 0.401 | 0.233 | 0.159 | -0.192 |
| Chores | -0.201 | 0.06 | -0.025 | 0.018 |
| Music | -0.455 | 0.05 | 0.158 | -0.017 |
| Resting | -0.433 | 0.376 | -0.145 | 0.032 |
| Nothing | -0.321 | 0.228 | 0.295 | 0.143 |
| Exercise | -0.176 | -0.238 | 0.615 | -0.216 |
| Hygiene | -0.096 | 0.026 | 0.325 | 0.077 |
| Reading | 0.2 | 0.107 | -0.181 | 0.219 |
| Shopping | 0.319 | -0.321 | 0.129 | 0.386 |
| Conversation | -0.189 | -0.215 | 0.076 | 0.333 |
| Phone-Call | 0.108 | -0.186 | -0.15 | 0.641 |
| Texting | -0.135 | 0.158 | 0.173 | 0.785 |
| Commuting | -0.176 | 0.041 | 0.323 | 0.236 |
| Computer | -0.195 | 0.009 | -0.093 | 0.098 |
| TV | -0.479 | -0.168 | -0.541 | 0.207 |
| Working | 0.603 | -0.158 | -0.066 | 0.346 |
| Other | -0.014 | 0.007 | -0.135 | 0.139 |

### Component Reliability

To further understand the robustness of the components from our analysis, we conducted a split half reliability for our sample. In this analysis we divided our data into two random samples and then examined how the factors generated in each half of the data related to each other. To compare the robustness of the solutions across a wide range of solutions we completed the component reliability analysis used PCAs with 3-, 4-, and 5-factors extracted (Figure S1, 2, S2). The mean correlation for the set of homologous pairs from each solution was calculated to determine which solution produced the most reproducible factors. The 4-factor solution produced the most reliable factors, with an average homologue similarity score of .953 (min. R_Hom._ = .93, max. R_Hom._ = .98) (Figure. 2).

**Figure 2.**
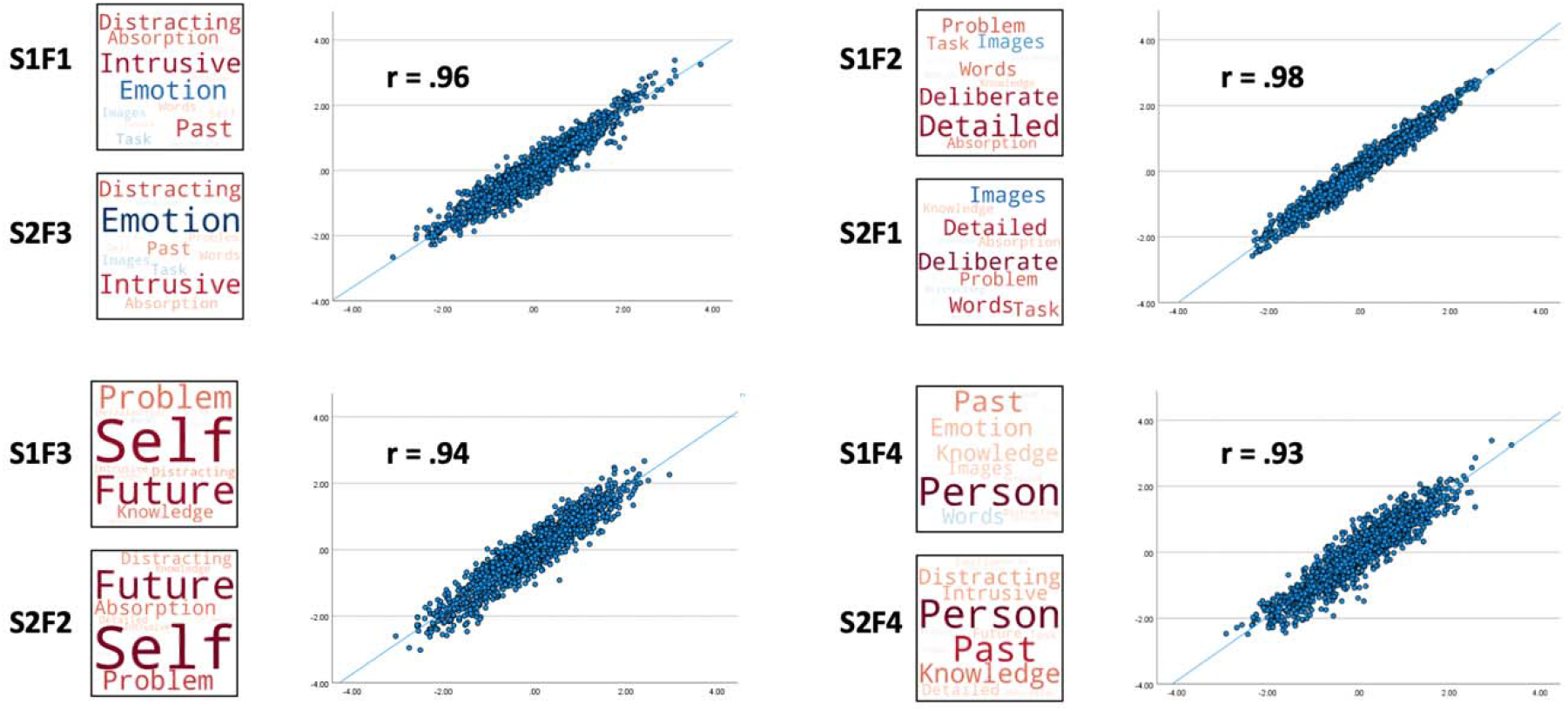
Component reliability analysis. Scatter plot of average homologue similarity. “S” indicates subset, and “F” indicates factor.

### The Influence of Socializing on Ongoing Thought

The first goal of our study was to replicate results from Mckeown et al. (2021). To do so, we compared the mean regression factor score for the episodic social cognition thought component across different types of social settings in physical and virtual environments (Figure 3). The episodic social cognition thought component varied significantly across both physical (*F*(2, 1403.16) = 20.75, *p* <.001) and virtual (*F*(2, 1403.32) = 18.83, *p* <.001) depending on the reported descriptions of the social environment. Based on the confidence intervals in Figure 3 it is clear that this pattern was prevalent when participants were around people and interacting with them either virtually or in person.

**Figure 3.**
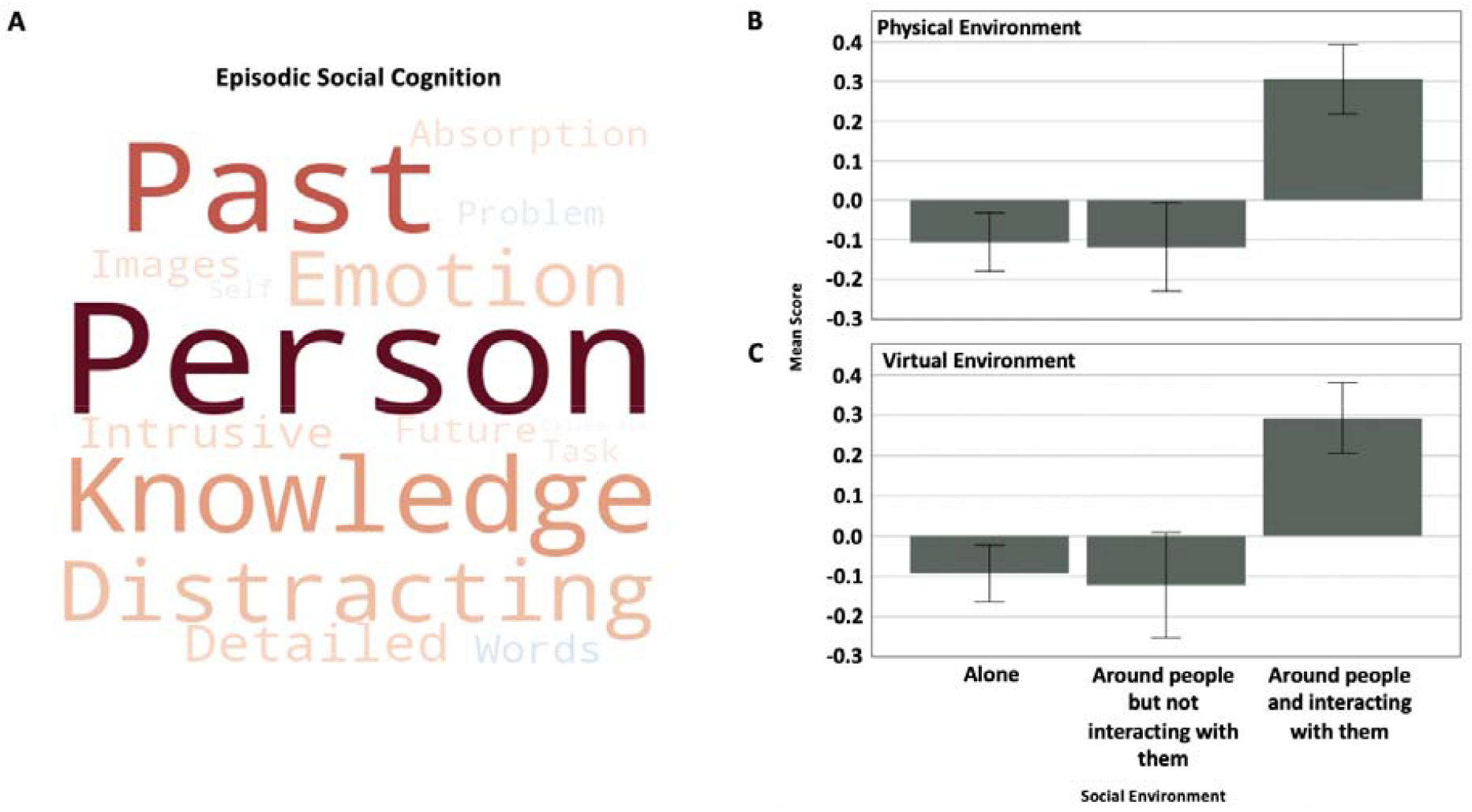
The influence of socializing on ongoing thought. (A) Episodic Social Cognition word cloud. Words represent experience-sampling items. Larger fonts are items with more importance and colour described direction (warm colours relate to positive loadings). (B) Bar chart comparing the mean multi-dimensional experience-sampling (MDES) scores when participants reported they were 1) alone, 2) physically around people but not interacting with them, and 3) physically around people and interacting with them. Error bars represent 99% CIs. (C) Bar chart comparing the mean MDES scores when participants reported they were 1) alone, 2) virtually around people but not interacting with them, and 3) virtually around people and interacting with them. Error bars represent 99% CIs.

### Thought-Activity Mappings

A second goal of our study was to extend research from the laboratory to examine whether associations between activities in the real-world and ongoing activities generalized beyond social interaction. In each case we found a significant association (Detailed Task Focus (*F*(17, 1412.78) = 11.70, *p* <.001), Negative Intrusive Distracting (*F*(17, 1388.15) = 3.86, *p* <.001), Future Problem-solving (*F*(17, 1395.30) = 4.88, *p* <.001), Episodic Social Cognition (*F*(17, 1399.13) = 4.53, *p* <.001)). To visualize these relationships, we generated a set of word clouds based on the loadings for each component for each activity for each component, and these are displayed in Figure 4. It can be seen that detailed task focus had high loadings when at work or doing homework, negative intrusive distracting thoughts had high loadings when resting or doing homework, future problem solving had high loadings when exercising and episodic social cognition had high loadings when texting, in conversation or on the phone.

**Figure 4.**
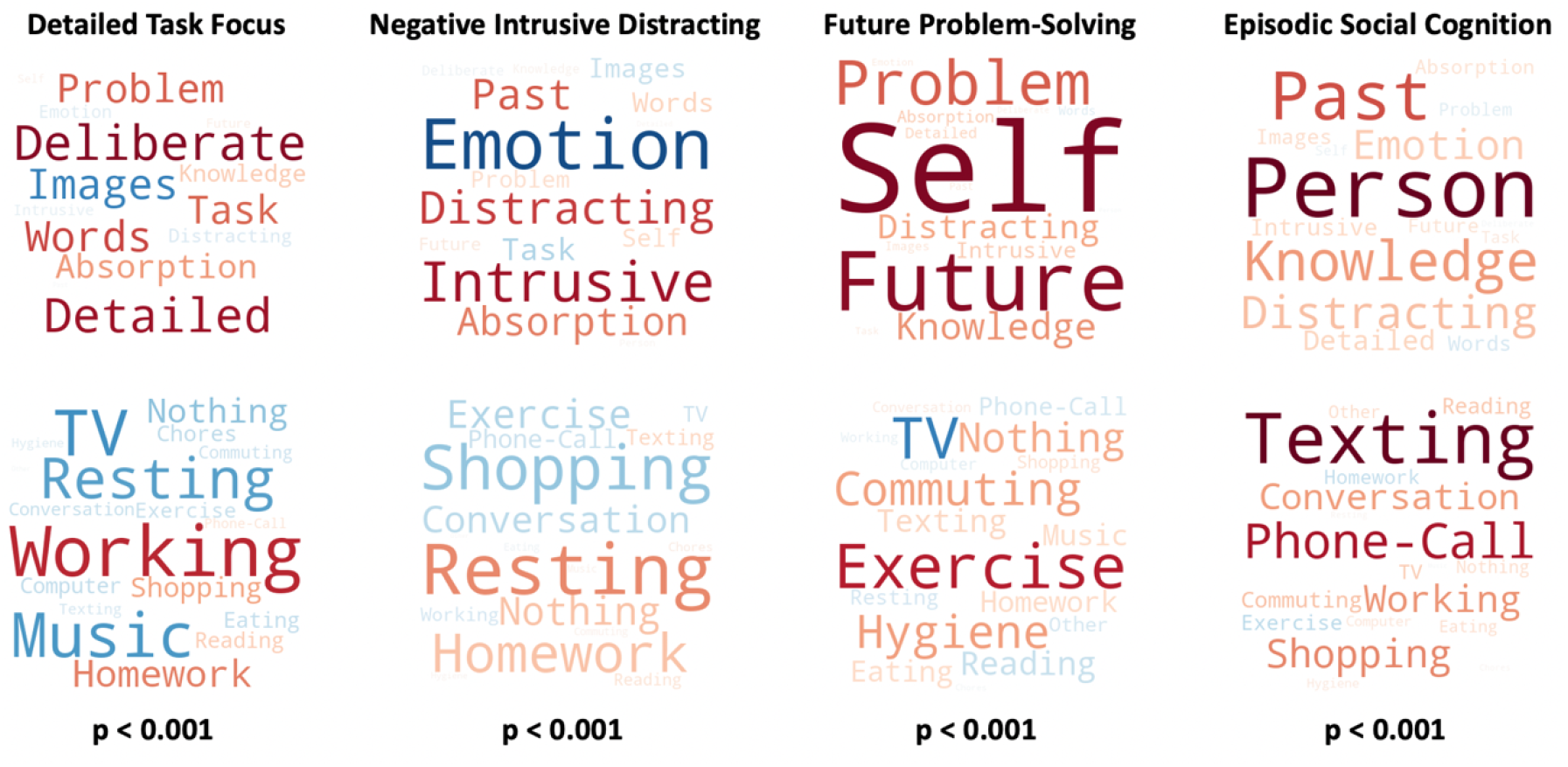
Thought and activity word cloud mappings. Words represent the PCA experience-sampling loadings, and LMM primary activity loadings. Larger fonts are items with more importance and colour described direction (warm colours relate to positive loadings). See Table 5 and Table 6 for specific component loadings.

To further visualize the relationship between ongoing thought and activities, LMM loadings were placed into three-dimensional spaces created by the four components. For simplicity in Figure 5 we generated a 3-dimensional space constructed from the episodic social cognition, future problem-solving and detailed task focus components (Figure 5A) and also included a 2-dimensional space was included to capture the relationship between episodic social cognition and negative intrusive distraction (Figure 5B).

**Figure 5.**
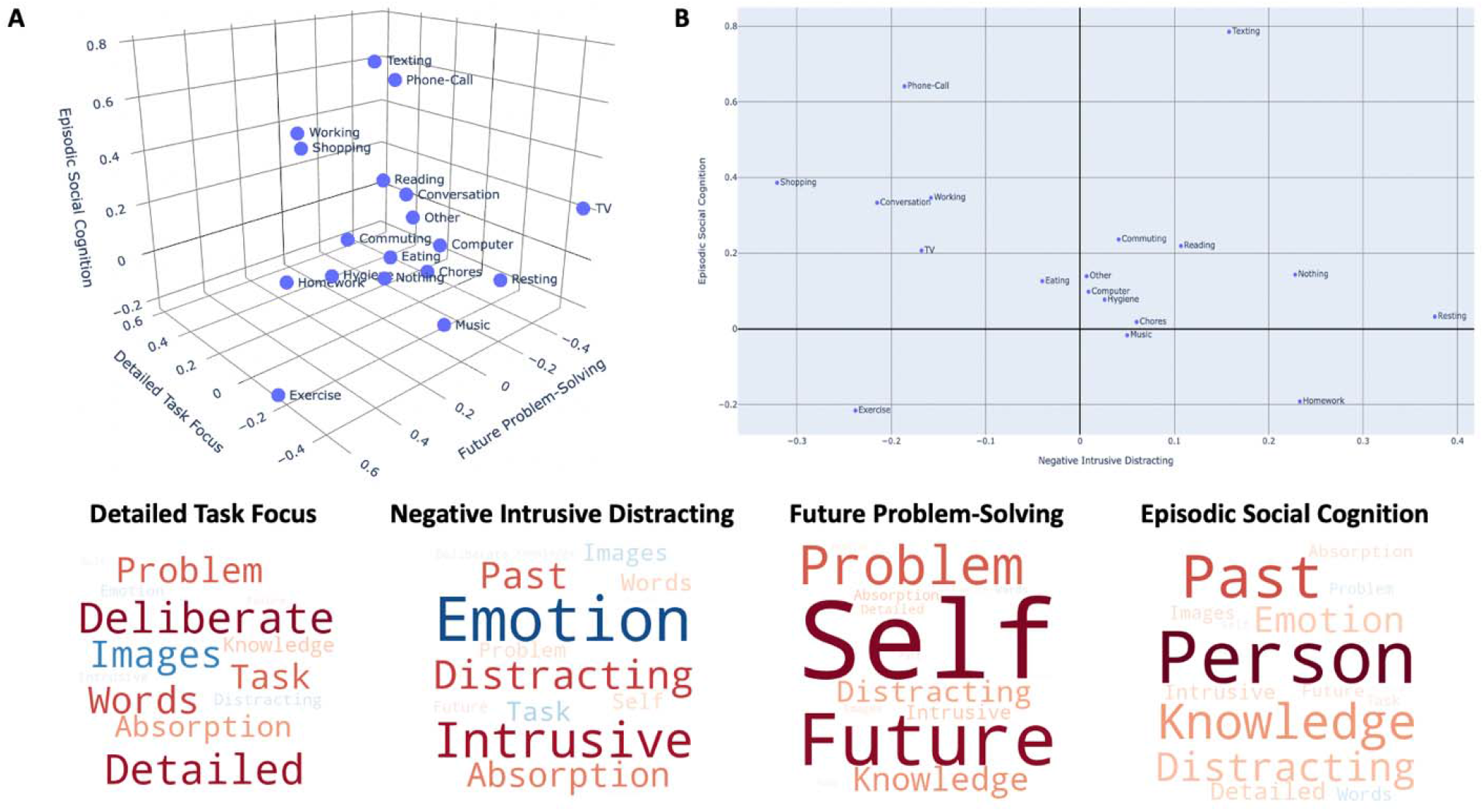
Mappings between MDES thought patterns and the activities in daily life. These data are presented in (A) 3- and (B) 2-dimensional spaces to provide an alternative way of visualizing the relationships between dimensions identified in our study.

### The Influence of Physical Location and Time of Day on Ongoing Thought

Having examined the links between activities and thought in daily life, we next turned to our two exploratory goals. First, we explored how physical location (inside or outside) related to the thoughts that people experienced. To do this, we conducted a LMM for each component score. Physical location was significant for the detailed task focus thought component (*F*(6, 1391.17) = 6.51, *p* <.001), which was higher when inside a workplace (Figure 6). No difference was found for the other components (Negative intrusive distracting (F(6, 1369.81) = 1.32, p = .22), future problem-solving (F(6, 1376.46) = .91, *p* = .484), and the episodic social cognition (F(6, 1381.34) = 1.38, *p* = .221). Next, we explored how the time of day when experience sampling occurred was reflected in differences in the patterns of ongoing thought that participants reported by conducting separate LMM for each component score. Time of day was significant for patterns of detailed task focus (*F*(5, 1408.69) = 4.27, *p* <.001) and episodic social cognition (*F*(5, 1394.20) = 4.20, *p* <.001), but not for the negative intrusive distracting (*F*(5, 1386.21) = .18, *p* = .969) and future problem-solving (*F*(5, 1391.13) = 1.92, *p =* .0881, Figure 7).

**Figure 6.**
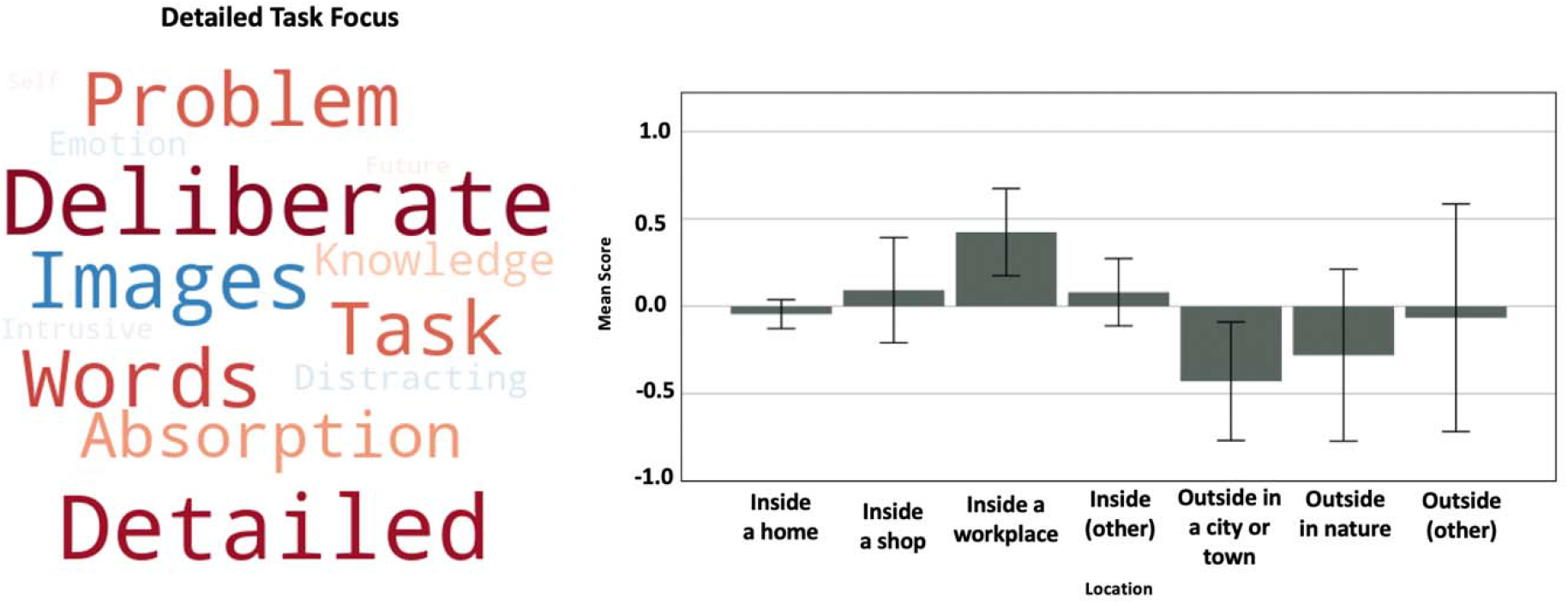
Results of an exploratory analysis comparing the relationships between the location of activities and ongoing thought patterns. We identified that patterns of detailed task focus were most prevalent in a work place, and least prevalent outside in either a city or a town. Error bars represent 99% CIs.

**Figure 7.**
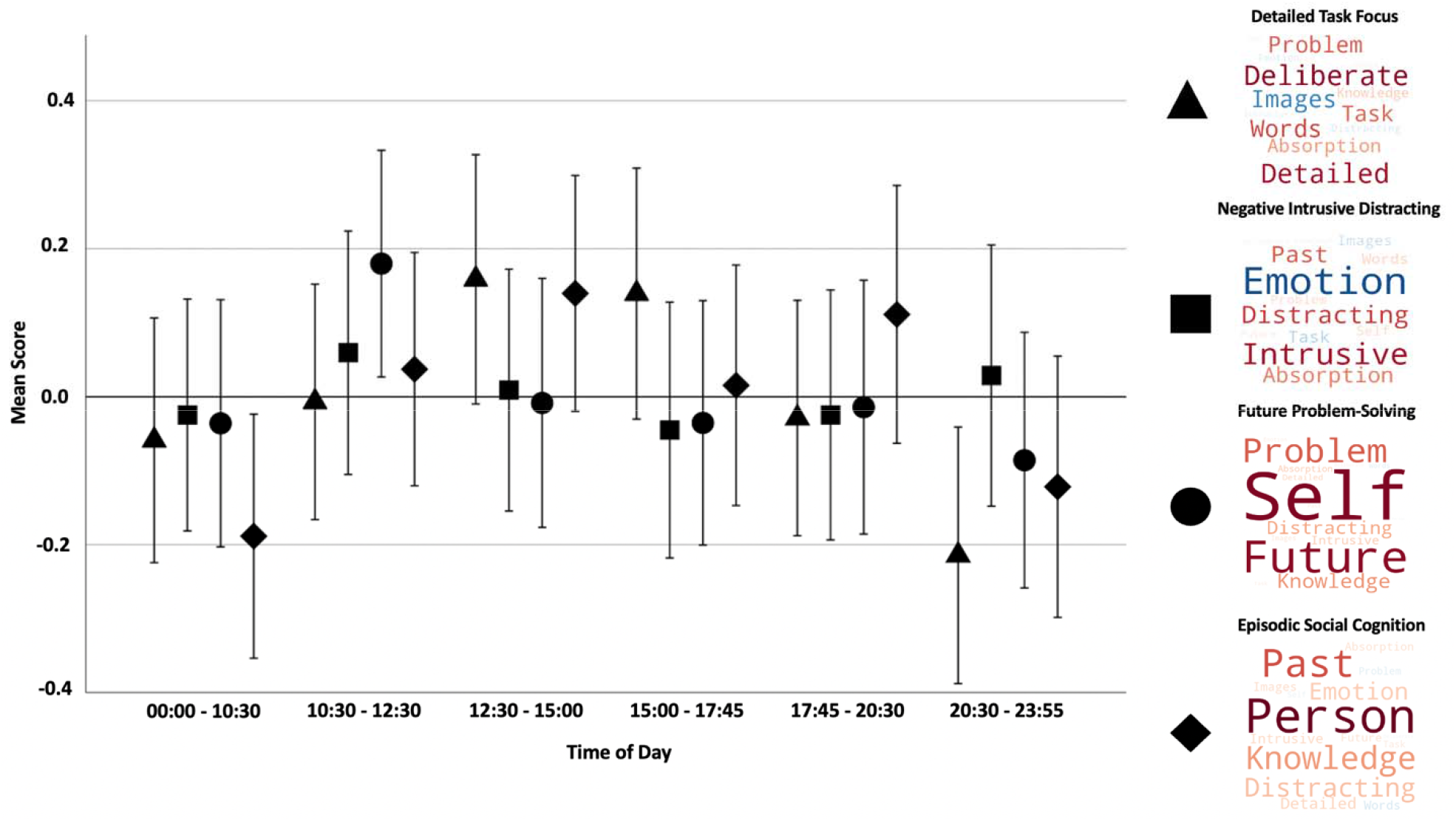
Results of an exploratory analysis examining the influence of time of day on ongoing thought patterns recorded by MDEs. The bar chart compares the mean score for experience-sampling responses across different time intervals: the morning (00:00:00 - 10:30), late morning (10:30 - 12:30), early afternoon (12:30 - 15:00), late afternoon (15:00 - 17:45), evening (17:45 - 20:30), and night (20:30 – 23:00). Error bars represent 99% CIs.

## Discussion

Our study set out to map patterns of ongoing thought and behaviour throughout real-world contexts. We hoped that measures of experience generated via multi-dimensional experience-sampling (MDES) would be able to differentiate the context in which the probes occurred, in particular the activities that people were performing. First, we sought to replicate the influence of socializing on patterns of ongoing thought found in Mckeown et al. (2021). Consistent with that study, we found that participants reported patterns of thought with episodic and social features when they were interacting with people in either a physical or a virtual manner.

We also examined if MDES can more broadly capture thinking patterns that reflect the sorts of activities participants performed in the real-world. A prior study had established that in the laboratory, MDES can capture patterns of thought that discriminate between the types of tasks that people performed (Smallwood et al., 2021; Konu et al., 2020). Consistent with this goal, we discovered general associations between the four ongoing thought patterns captured by MDES and the everyday activities people were performing. Detailed task focus thought patterns were most prevalent when people are working and doing homework. This pattern of thought in the lab is known to emerge consistently when participants perform tasks which demand executive control including working memory or task switching (Sormaz et al., 2018, Turnbull et al., 2020; Konu et al., 2021). Future problem-solving thought patterns were present during activities like exercise, commuting, and doing nothing. In the lab this style of thinking emerges when task demands are lower (Turnbull et al., 2020; Ruby et al., 2013), and can be associated with an individual’s generating patterns of personal goals with greater details (Medea et al., 2018). Negative intrusive distracting thought patterns were present when resting, doing nothing, and homework. Finally, consistent with an association with social cognition (Mckeown et al., 2021; Konu et al., 2021) patterns of episodic social thought were present in activities which likely involve other people including conversations, texting, and shopping. Intriguingly, task studies have shown that this pattern of thought emerges when people make decision on a familiar other (Konu et al., 2021) and brain imaging studies have shown that this pattern of thought is linked to activity in the medial prefrontal cortex (Konu et al., 2020).

We also had two more exploratory goals. We first examined how the physical location (inside or outside) was related to the thoughts that people experienced. This analysis identified that detailed task focus thought patterns were present when participants were inside a workplace and absent when they were inside a home, inside a shop, inside (other), outside in a city or town, outside in nature, or outside (other). Location was not significant for negative intrusive distracting, future problem-solving or episodic social cognition thought patterns. Second, we examined how the time of day was reflected in participants responses to the MDES probes. This exploratory analysis found that patterns of detailed task focus were more likely to be reported in the middle of the day, and less likely at night. Similarly, episodic social cognition thought patterns were least present in the early morning, and most common in the early afternoon. Note that these exploratory analyses should not be taken to indicate direct consequences of location or time of day on ongoing experience. Instead, these effects are likely to indicate that the activities themselves are more likely to occur in specific locations or at particular times of the day. Disentangling the specific variables which drive these associations is likely to be important in future studies.

In conclusion, our results suggest that patterns of thinking in the real-world indirectly reflects the situation in which experiences emerge. Our study suggests that ongoing activities are likely to be important in the types of thoughts a person has, and that other factors such as location or time of the day may contribution to this phenomenon less directly. This indicates that MDES is able to differentiate between the different situations that people are in within daily life. This highlights the value of MDES as a tool for understanding cognition from an ecological perspective, particularly because this technique can also be used in more controlled settings, such as work that uses this technique in conjunction with brain imaging to reveal the neural correlates of different thought patterns (Turnbull et al., 2019, 2020; Konu et al., 2020). In this way MDES may be a useful tool for bridging the gap between more controlled laboratory settings, where specific features of cognition can be directly manipulated by the experimenter, and more realistic situations in daily life. In this way experience sampling in daily life, and in particular techniques like MDES may be an important next step in building accounts of cognition that are more ecologically valid (Kingstone et al., 2003).

Although our data establishes that MDES is a useful tool for mapping cognition in daily life, our data also raise a number of open question that future research could address. First, data collection began during a COVID-19 lockdown, reducing the types of activities participants could self-select, and potentially biasing the patterns of thoughts that our study identified. Thus, while our study clearly shows the utility of MDES and experience sampling (ES), there may be types of activities, and therefore, patterns of experience, that would be captured by ES in daily life outside of a lockdown situation. Additionally, notification response rate and timing varied across participants, which could relate to participant motivation or possibly activity enjoyment or value. Specifically, participants may have been less likely to immediately respond, or to respond at all, to a notification during particularly enjoyable activities. Furthermore, study participants were students enrolled in designated first- and second-year psychology courses, with an average of 21 years. Participant age and occupation is likely to be an important factor in regard to the types of activities self-selected, and thus, the thought components produced in our study may be less generalizable to a broader more representative sample. Lastly, potential thought components are limited by the choice of MDES probes offered to the participants in our study. Although the items we used can dissociate the links between activity and thoughts, with more accurate question this capacity could be improved. For example, during the analysis process, it was noted that the detailed task focus component was negatively anchored by music and TV. Although images may be a useful characteristic of watching TV, it is less useful as a way to characterize their thoughts while they listen to music. Future studies using MDES, therefore, could benefit from breaking the modality probe into three questions, giving participants the opportunity to describe their experience in terms of images, words, and/or sounds.

Finally, we close by noting that by sampling thinking in the daily life our results are likely to depend in a complex way on how people select the activities they perform in daily life. Presumably, individuals have a degree of choice about the tasks they perform in daily life that is absent for many laboratory studies (Kahneman et al., 2004; Smallwood et al., 2021). For example, it is likely that more sociable individuals spend more time engaged in forms of social cognition, more athletic individuals engage in exercise more often, and more studious individuals spend more time on their homework. Accordingly, our study suggests that when sampling cognition in naturally occurring situations, temperament or expertise in a specific domain may be indirectly related to the thought patterns they experience, as people may perform activities that they enjoy or are good at when outside of a laboratory setting. This ability to choose the activities in our daily life may be a primary reason why thought patterns in the lab do not always generalise to the real world (Kane et al., 2017; Ho et al., 2020).

